# High resolution optical spectroscopy for the evaluation of cannabidiol efficiency as a radiation therapy support of peripheral nervous system tumors

**DOI:** 10.1101/2023.12.11.571087

**Authors:** Karolina Chrabąszcz, Katarzyna Pogoda, Klaudia Cieżak, Agnieszka Panek, Wojciech M. Kwiatek

## Abstract

An increasing number of scientific papers discuss the promising therapeutic potential of cannabidiol (CBD) not only for the treatment of cancer, but also for asthma and neurodegenerative disorders. This happens mainly due to its proven anticancer, anti-inflammatory, and antioxidant properties. In the field of cancer research, the use of CBD has already been investigated on malignant tumors of the central nervous system, like gliomas. So far, CBD has not yet been explored in the therapy of peripheral nervous system (PNS) tumors. Peripheral nerves reside outside the central nervous system, therefore peripheral nerve tumors can occur anywhere in the body. When the tumor develops within large blood vessels, spinal nerves or involves more than one peripheral nerve, radiotherapy is recommended. Due to high doses of ionizing radiation, complications such as dizziness, damage to adjacent nerves, or malignancy of the lesion may occur. Therefore, it is important to develop a treatment scheme that efficiently reduces tumor volume while maintaining the normal functions of the surrounding cells and decrease the side effects. Herein, we proposed to combine hyperspectral imaging using Raman and FTIR spectroscopy and AFM-IR technique as a novel approach to monitor the therapeutic efficacy of CBD. Performed studies reviled the dual effect of CBD, that protects normal cells from ionizing radiation and increases its toxicity in cancer cells.

## 1. Introduction

Cancer remains in the top three of the deadliest diseases worldwide, moreover, it is estimated that in 2060 it will be the leading cause of death.^1^ Although innovative anticancer therapies are under investigation; radiotherapy is still one of the most used treatment approach.^2^ Radiation therapy is especially beneficial if the cancer affects smaller areas, is located adjacent to blood vessels or any essential structure which may be damaged during surgery. Also for patients with incurable cancers, radiotherapy allows for relieve of the symptoms (palliative radiotherapy).^3^ Since radiation therapy uses high doses of ionizing radiation, it is important to look for a treatment that allows for the efficient reduction of the tumor volume while maintaining the normal functions of the surrounding cells and decrease of the side effects.

Nowadays, various natural therapies gain more attention as adjunctive cancer treatment or anti-side effect substances.^4^ Cannabidiol (CBD) is non-psychoactive phytocannabinoid known for its proven anti-cancer, anti-inflammatory, antioxidant, and neuroprotective properties.^5, 6^ Due to the range of CBD advantages it is investigated in preclinical models as an adjunctive substance for cancer treatment, chronic pain, asthma, and neurodegenerative diseases.^7,8,9^ Moreover, Food and Drug Administration (FDA) approved a cannabidiol-based drug (Epidiolex®) for the treatment of Lennox-Gastaut or Dravet syndrome with episodes of epilepsy in children below 4 years old.^10, 11, 12^

In the field of cancer research, the use of CBD has already been investigated on malignant tumors of the central nervous system, like gliomas, however its effectiveness in the peripheral nervous system (PNS) tumors so far remains unknown.^13, 14^ Due to the fact that peripheral nerves branches outward from the spinal cord and brain, cancers which arise from PNS may appear anywhere in the body.^15^ Neoplasms that affect peripheral nerves can form benign and malignant peripheral nerve sheath tumors (MPNST) which originate mainly from Schwann cells.^16^ MPNST belongs to an aggressive soft tissue sarcomas, with limited therapeutic options and poor prognosis.^17^ So far, the only known definitive therapy is the complete resection with wide margins of malignant lesions.^18^ Approximately 30-50% of the people who suffer from the genetic disorder called neurofibromatosis-1 (NF-1) develop plexiform neurofibromatosis which tends to transform into MPNST.^19, 20^ These lesions are often difficult to diagnose because they resemble other subcutaneous tumors such as lipomas or multiform adenomas. Lack of early diagnosis leads to the growth of these tumors, inducing compression on the surrounding blood vessels and nerves, leading to their damage.^21^ For MPNST which develops within cranial nerves such as the vestibular nerve, uncontrolled growth may cause hearing loss and balance disorders.^22, 23^ In that case, when the surgery is not applicable due to the possibility of damaging important adjacent tissues, stereotactic radiosurgery is recommended.^24, 25^ This therapy involves the administration of one or several (<5 fractions) high doses of ionizing radiation (5-16 Gy), which often cause complications such as dizziness, damage to adjacent nerves, including the auditory nerve, or malignancy of the lesion.^26, 27^ For that reason, it is essential to consider a substance that will increase the toxicity of ionizing radiation in tumors and at the same time protect surrounding cells. Due to the fact that CBD exhibits both anti-cancer and neuroprotective effects it may have the potential to fulfill these criteria as a novel adjunct agent for radiation therapy.

Focusing on CBD applications on various cancerous and normal cell lines as well as mice models, each study consists of multiple quantitative methods to determine changes in protein and lipid fractions (western blot, liquid chromatography), DNA (RT-PCR), or cell morphology (fluorescent staining).^28,29,30^ However, these methods require the use of expensive chemical reagents during complicated multi-stage processes which usually damage the studied sample. Reaching the need for obtaining rapid outcomes from sample analysis the various spectroscopic techniques of Raman (RS) and Infrared (IR) spectroscopy show potential as objective, label-free, and fast analytical methods.^31, 32^ Spectroscopic techniques do not require complex sample preparation protocols and data collection (usually several thousands of spectra) takes place in a relatively short time (up to several minutes). Acquired chemical images allow not only for monitoring the total chemical composition of a studied sample but also the distribution of particular molecules even at the subcellular level as well as give information about biochemical or morphological changes related to the disease or therapy process, even enabling the assessment of the radiation effects.^33,34,35,36,37^ Moreover, the combination of atomic force microscope (AFM) with IR technique (AFM-IR), allows for the investigation of biochemical changes at the nanoscale level and with the accurate representation of the topography of the studied sample.^38, 39^,

Herein, for the first time, we propose the use of hyperspectral imaging and AFM-IR technique as a novel approach to monitor the therapeutic effectiveness of cannabidiol in the treatment of peripheral nervous system tumors.

## 2. Materials and Methods

### 2.1 Cannabidiol

Cannabidiol (CBD) solution 1.0 mg/mL in methanol was purchased from Merc. The solution was initially dissolved in methanol to obtain the 1000μM concentration and stored at −20°C. It was further diluted in a cultured medium to the desired concentrations for cell studies. The dilutions were made considering the methanol concentration below 0,001%, to exclude methanol toxicity on cells.

### 2.2 Cell culture

Studies were conducted on two cell lines purchased from ATCC: Human Schwann cells isolated from the peripheral nerve trunk (normal, hTERT NF1 ipnNF95.11c) and human MPNST derived from lung metastasis (cancer, sNF02.2). Cells were cultured according to ATCC protocol in DMEM medium, supplemented with 10% of FBS (fetal bovine serum), 100 U/ml penicillin–streptomycin-neomycin solution. MPNST cell line was additionally supplemented with L-glutamine. The cells were grown in75 cm^3^ flasks at 37°C in an atmosphere of 5% CO_2_.

For spectroscopic studies cells were seeded on calcium fluoride windows (Crystran Ltd., UK) inside 12-well plates and kept in an incubator at 5% CO_2_ and 37°C for 24 h to promote adhesion and growth. For each cell line, 2 sets of samples were prepared: (I) incubated with CBD for 24h, (II) incubated with CBD for 24h, and then exposed to 10 Gy X-ray radiation (Fig. 1). For each group also the control samples (0μM CBD) was prepared.

**Figure 1.**
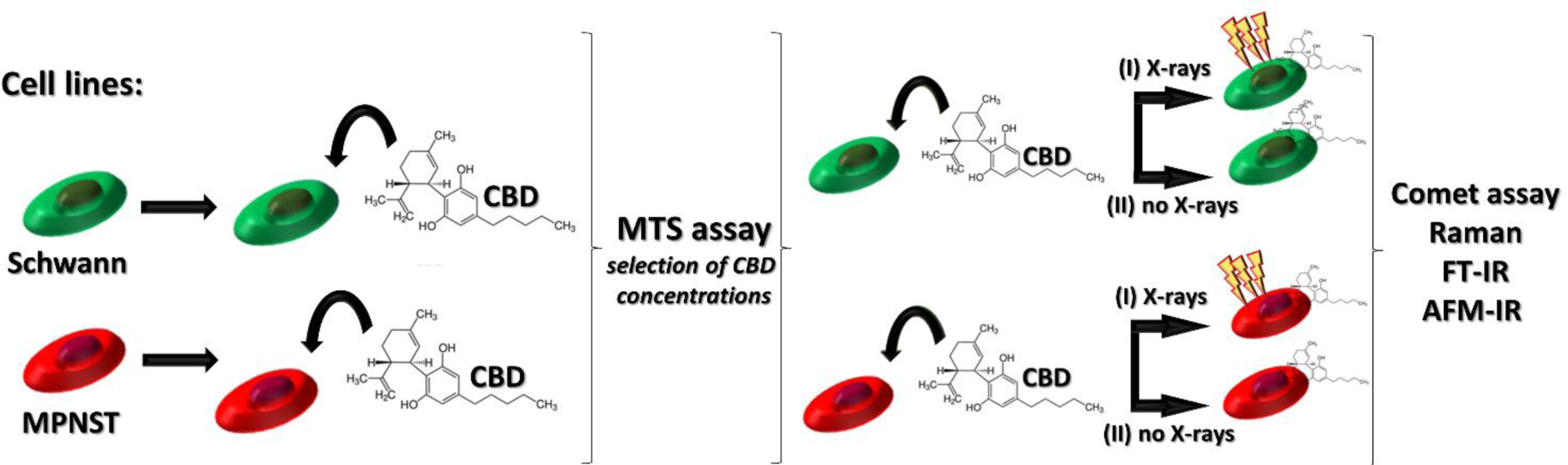
Experimental workflow.

### 2.3 MTS assay

To investigate CBD influence on the cells viability and metabolic activity the cells were tested using CellTiter 96® AQueous One Solution Cell Proliferation Assay (Promega) with tetrazolium compound. Cells were seeded in 24-well plates at a density of 15, 000 /well and 24h after seeding cells were serum-starved for 2h in DPBD with Ca^2+^ and Mg^2+^. Next, cells were treated with a selected concentration of CBD. Since the MTS assay is performed on live cells, particular MTS read-outs after 24h, 48h, and 72h of incubation with CBD were carried out on a single 24-well plate, and not at three separate plates, for every time point. The scheme of workflow is presented in Figure 1. CellTiter 96® AQueous One Solution reagent was added directly to the culture wells and incubated for 1 hour. Then the absorbance of formazan was recorded at 490 nm with a Spark 10 M (Tecan) multimode microplate reader. The absorbance values obtained with the untreated cells (control) were used for data normalization. Assays were conducted in triplicate. For spectroscopic measurements 24h time point was selected.

### 2.4 Irradiation procedure

For irradiation studies cells were seeded on calcium fluoride windows (Crystran Ltd., UK) inside 12-well plates and kept in an incubator at 5% CO_2_ and 37°C for 24 h to promote adhesion and growth. The cell confluence after that time was ca. 70%. For each cell line, 2 sets of samples were prepared as presented in Figure 1. The irradiation with a single fraction of X-rays at a dose rate of 2.1 Gy min^-1^ was performed using an MG325 (250 kV, 10 mA) X-ray tube (YXLON, Hamburg, Germany). For this experiment, 10Gy exposure dose was selected as commonly used for MPNST radiotherapy. To maintain the same experimental condition, the set of non-irradiated samples were kept out of the incubator (Fig. 1I) during the irradiation of second set of samples (Fig. 1II).

### 2.5 Comet assay

Comet assay was applied to analyze the genotoxic effect of CBD and CBD in combination with X-ray radiation. DNA damage levels were carried out directly after 24 h incubation with CBD and after 24h incubation with CBD and irradiation using the alkaline version of the comet assay. Suspension (50 µl) of Schwann and MPNST cells (3000-4000 per 1 ml) have been embedded in 150 µl of agarose on a microscope slide. Cells were lysed for 1h by 1% Triton X- 100 in pH> 13. Alkaline electrophoresis (30 V, and 300 mA) was then carried out for 30 min at 4°C. Ethidium bromide (17 mg/ml) was used for cells staining. Details of the applied protocol can be found in Refs. ^40, 41^ Epifluorescence microscope Olympus BX-50 connected with a CCD camera (excitation filter 515–560 nm, barrier filter from 590 nm) was used for cellular DNA visualization. Semi - automated image analysis system Komet 3.0 software (Kinetic Imaging Co., Liverpool, UK) was applied for the analysis. The extent of DNA damage was determined by the T-DNA parameter – tail DNA (DNA percentage in the comet tail). Two independent experimental replicates were performed for each aliquot: from 200 to 500 cells were analyzed for each data point (2 slides per each CBD concentration, without and with 10Gy X-ray dose, 100–250 cells from each slide). Pearson’s r correlation coefficient for data presented from comet assay were calculated from linear fitting.

### 2.6 Spectroscopic measurements

In order to perform spectroscopic studies, both cell lines were seeded at low density (30,000 cells/well) on CaF_2_ optical windows (Crystran Ltd) in 12-well cell culture plates. Each seeding of cell lines were prepared in duplicates, according to Fig. 1. 24 h after seeding, cells were serum-starved for 2 h in a serum free medium and then treated with the particular CBD concentrations for 24h or left untreated as control samples (0μM CBD) and irradiated with X-rays as described in section 2.4. The next step was rinsing twice with PBS for 2 min and fixing with 4% paraformaldehyde (PFA) in phosphate-buffered saline (PBS) for 20 min. Then, all samples were washed with PBS (3 times for 2 min) to remove PFA residues. For Raman measurements, samples were left in PBS solution. Before FT-IR and AFM-IR measurements samples were washed in PBS/water solutions (9:1, 8:2, 7:3; 6:4, 5:5, 4:6, 3:7,2:8,1:9, for 2 min), and ultrapure water then dried under the gentle stream of nitrogen. All solutions were prepared using ultrapure water (Direct-Q 3 UV, Millipore, USA).

#### 2.6.1 Raman microspectroscopy

Raman images were recorded using a Renishaw InVia Raman spectrometer equipped with an optical confocal microscope, an air-cooled solid-state laser emitting at 532 nm, and a CCD detector cooled to −70°C. An immersive Leica N PLAN EPI (60x, NA 0.75) objective was used. Raman maps were collected from the whole cell area for 15 cells per condition with 1µm step size, integration time of 1 s for 1 accumulation, and a spectral resolution of ca. 1.5 cm^-1^. The spectrometer calibration was performed with an internal silicon plate.

#### 2.6.2 FT-IR microspectroscopy

FT-IR images were collected using a HYPERION 3000 FT-IR microscope with 36x magnification objective, coupled with Vertex 70v spectrometer (Bruker, Ettlingen, Germany) operating in transmission mode. Hyperspectral images were recorded by the FPA detector of 64 × 64 pixels and projected pixel size 1.1 μm × 1.1 μm. Maps were collected from 15 cells for each condition (the spectral range from 900 cm^−1^ to 3800 cm^−1^, spectral resolution of 4 cm^−1^ and 256 scans per spectrum).

#### 2.6.3 AFM-IR nanospectroscopy

AFM-IR spectra (10 cells per condition) and exemplary maps were collected in contact mode using the NanoIR2 spectrometer (Anasys Instrument, Santa Barbara, California) with silicon gold-coated PR-EX-nIR2 probes (30 nm tip diameter, 13 ± 4 kHz resonance frequency, (Anasys Instruments, USA). Contact resonances were selected using a 180 kHz search location and a 50 kHz half-width Gaussian filter. For spectra collection, an optical parametric oscillator (OPO) laser was used as an infrared excitation source in the range of 3500-2800 cm^-1^ and 1800- 900 cm^-1^ with the 4 cm^-1^ spectral resolution. Spectra were collected from 7 points that were placed diagonally across each cell and condition by co-averaging 256 pulses of excitation. For the AFM topography images collection cantilever scan rate was set to 0.05 Hz and a spatial resolution of the 20×20nm images was 500 x 500 pixels. IR maps corresponding to topography images were collected for selected wavenumbers with a multichip tunable quantum cascade laser as an infrared source (QCL; MIRcat-QT Daylight Solutions).

### 2.7 Data analysis

Raman images were analyzed using WiRE (ver. 5.3, Renishaw, United Kingdom) software. Preprocessing included cosmic ray removal, noise filtering, baseline correction, and min-max normalization on all spectra. Then the hierarchical cluster analysis (HCA) was performed to differentiate the whole cell area and particular cellular components such as cytoplasm and nuclei based on their spectral profile. Euclidean distance and Ward’s algorithm were used to calculate spectral distances and the individual clusters. For FT-IR data preprocessing OPUS (ver. 7.5, Bruker Optics, Germany) software was used. Mean averaged spectra from every single cell were extracted and min-max normalization in the amide I region was performed to avoid differentiation to the sample thickness. Then the second derivative IR spectra were calculated with 13 smoothing points according to the Savitzky–Golay protocol. ^35, 42, 43^

The areas under the selected Raman and FT-IR bands (integral intensities) were calculated using OPUS (ver. 7.5, Bruker Optics, Germany). Based on the values of integral intensities box plots were constructed to obtain information about biochemical changes within cells. The variance was performed using the statistical model (ANOVA) in the OriginPro software. Tukey’s test was employed to compute significance values p.

The AFM-IR spectra were analyzed using Analysis Studio (ver. 3.14) software. Spectra were smoothed according to the Savitzky-Golay protocol with 3rd-order polynomial and 5 data points, normalized (min-max normalization) and presented as second derivative IR spectra.^44,45,46^ The 3D AFM and AFM-IR images were prepared with Mountainsmap software (ver. 7.3, Digital Surf, France)

## 3. Results and discussion

### 3.1 Cyto- and genotoxicity effects of CBD itself and CBD with the combination of ionizing radiation

In order to investigate the CBD influence on Schwann and MPNST cells viability, various CBD concentrations were tested (Fig. 3A). Interestingly, after 24h of incubation, increased viability of Schwann cells were observed, in the range of 0.01 μM to 5μM CBD concentration, compared to untreated, control cells. Surprisingly, CBD treatment did not affect MPNST cell viability, despite its proven anticancer effect. ^5,47^ For further investigations 0.025 μM, 0.1 μM, 0.5 μM, 3 μM and 9 μM concentrations were selected.

**Figure 2.**
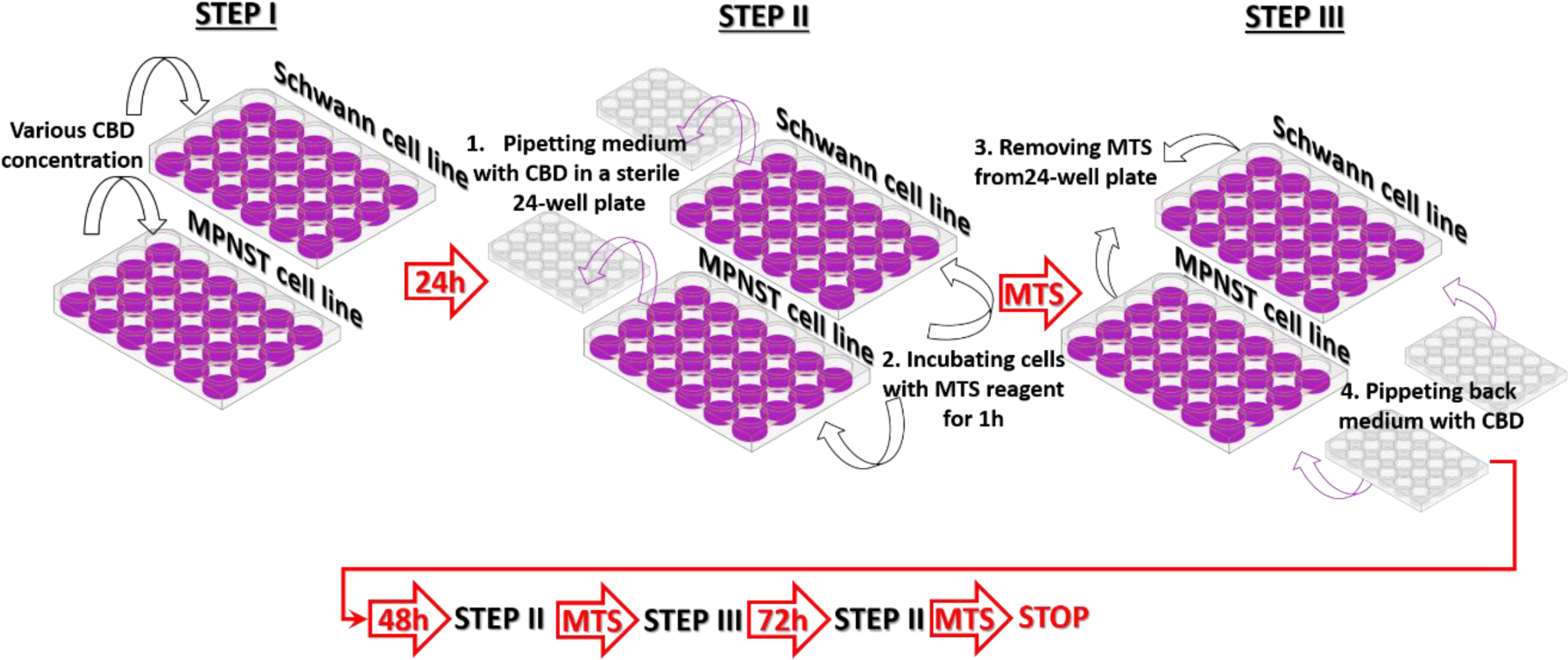
Schematic workflow of MTS assay cytotoxicity test.

**Figure 3.**
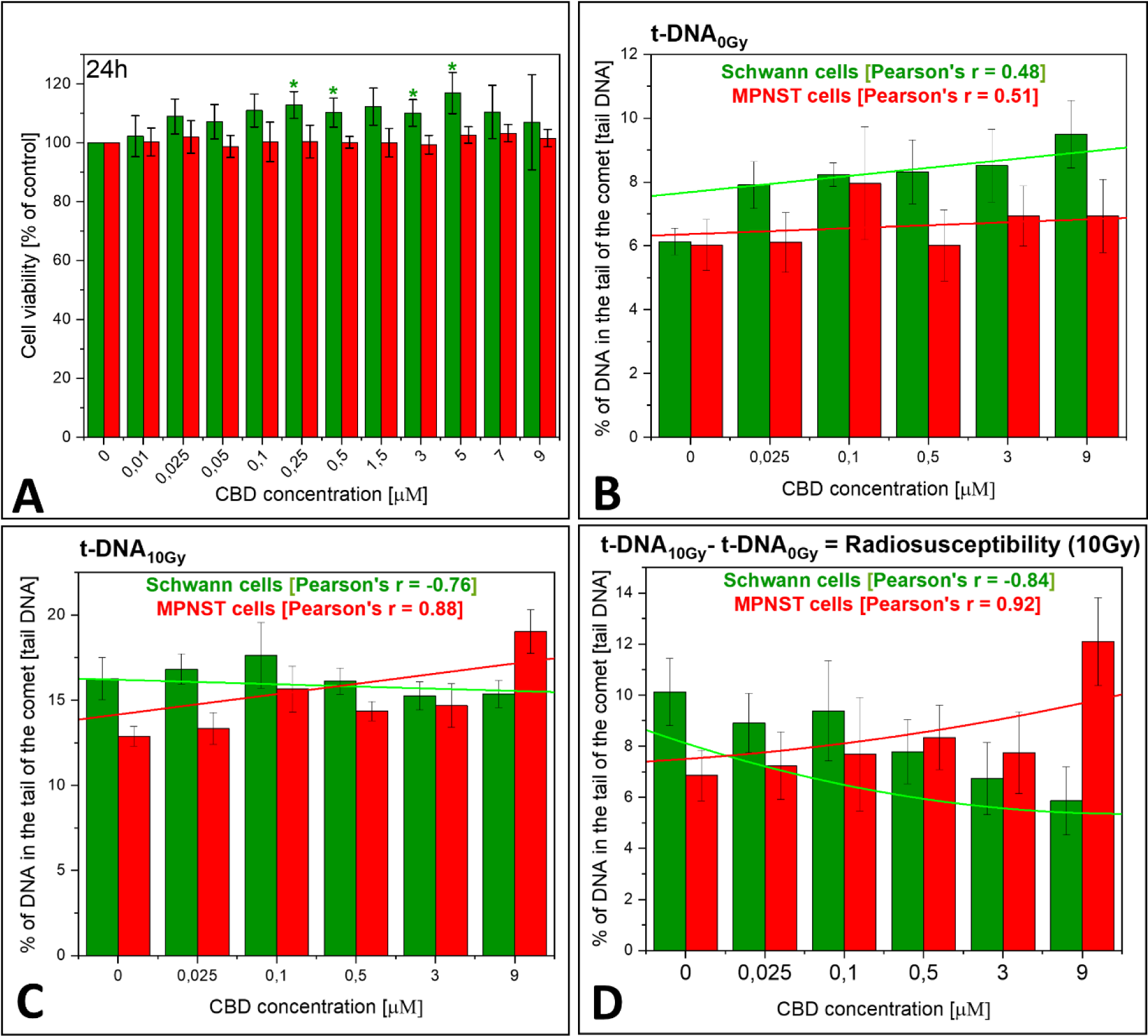
Results of MTS test for twelve CBD concentrations. (A) The statistical significance was calculated in contrary to untreated cells, with ANOVA test (*p<0.05). Results of comet assay (B) for selected CBD concentrations, (C) for selected CBD concentrations and 10Gy irradiation dose, (D) radiosuscebility for 10Gy. Pearson’s correlation coefficient was calculated from linear fitting.

Due to the fact, that radiotherapy (ionizing radiation) induces single and double-strand DNA breaks, it was essential to firstly estimate the possible influence of CBD itself on DNA structure before X-ray irradiation was applied. For that purpose, comet assay was performed on both cell lines. Based on electrophoresis and fluorescence staining it was possible to observe the damaged DNA, presented as comet “tail” separated from the intact DNA “head”. From this test, the t-DNA_0Gy_ (tail DNA) values were calculated for both cell lines incubated with selected CBD concentrations, without irradiation procedure (0Gy) (Fig. 3B). The Pearson correlation coefficient statistically estimate the strength of a linear relationship between paired data, denoted by r and constrained as −1 ≤ r ≥ 1. R values were calculated form the linear fitting and for t-DNA_0Gy_ they amount 0.48 and 0.51 for Schwann and MPNST cells, respectively. Such values include in the 0.4-0.59 range (positive moderate correlation), what indicate some relationship between the CBD concentration and level of DNA damage for both cell lines, however the indirect relationship is not strong. Therefore, CBD itself did not reveal any toxic effect on DNA for Schwann and MPNST cell lines (Fig 3. B). On the contrary, the t-DNA_10Gy_ established for cells treated with CBD and irradiated with 10Gy exposure dose display differences in the cells response to ionizing radiation, what depend on cell lines (Fig 3. C). For Schwann cell line strong negative correlation appears (−0.76 in the range from −1 to −0.70) what indicate that the highest CBD concentration, than less DNA damage caused by ionizing radiation occurs. Interestingly, an opposing strong positive correlation (0.88 in the range from 0.7 to 1) is visible for MPNST, therefore the increase in CBD concentration enhances the toxic effect of radiotherapy, what manifests in increase of damaged DNA observed in comet “tail” Based on t-DNA_0Gy_ and t-DNA_10Gy_ values it was possible to evaluate the relative susceptibility of cells to ionizing radiation. Radiosusceptibility was calculated as the difference in the levels of DNA damage immediately after (t-DNA_10Gy_) and before (t-DNA_0Gy_) radiation (Figure 3D). Surprisingly, with an increase in the CBD concentration, the contrary cell response to 10Gy exposure dose can be seen. Both cell lines presents strong correlation, however for Schwann the negative correlation value (−0.84) displays the decrease in radiosusceptibility with an increased CBD concentrations where for MPNST high positive correlation (0.92) indicate increase in radiosusceptibility as the CBD concentration increase. Therefore, with an increase CBD concentration, under the influence of 10Gy irradiation exposure dose the DNA of Schwann cells is less damaged and become more resistant to ionizing radiation, whereas for MPNST it enhances DNA destruction and increase vulnerability to radiation.

### 3.2 Spectroscopic signature of CBD and CBD combined with radiation influence on Schwann and MPNST cells

Since hyperspectral imaging, as Raman and FT-IR imaging, allow for simultaneous acquisition of biochemical information (spectra) with its distribution (chemical maps) in a micrometric scale, spectroscopic techniques can shed a light on detailed biochemical changes occurring at the subcellular level, which reveal the CBDs’ mechanism of action as a potential adjuvant agent for radiotherapy.

Hyperspectral imaging generates thousands of spectra per single cell, for that reason the data analysis based on clustering is useful for constraining the number of spectra in collected datasets. Cluster analysis (CA) allows for the spectral grouping based on their profile similarity. In case of acquired cells hyperspectral images, with CA is possible to differentiate the area occupied by whole cell as well as distinguish sub-cellular component as nuclei, cytoplasm, endoplasmic reticulum and lipid droplets.^48, 49^ Because the purpose of this work was to establish the overall cell response to CBD and radiation, CA was used for selection mean averaged Raman and FT-IR spectra, which represents biochemical composition of particular cells.

Mean Raman spectra obtained from the collected maps of single cells display subtle changes, mainly in the high-wavenumber range associated with the alterations in the intensity for the 2850 cm^-1^ band related to CH_2_ vibration from long-chain lipids (Fig. 4, *top panel*). Because subtle changes are hardly noticeable based on spectral profile, to emphasize biochemical changes occurring under the influence of CBD and its combination with ionizing radiation integral intensity of selected bands were calculated and presented as box plots (Fig. 4, *bottom panel*).

**Figure 4.**
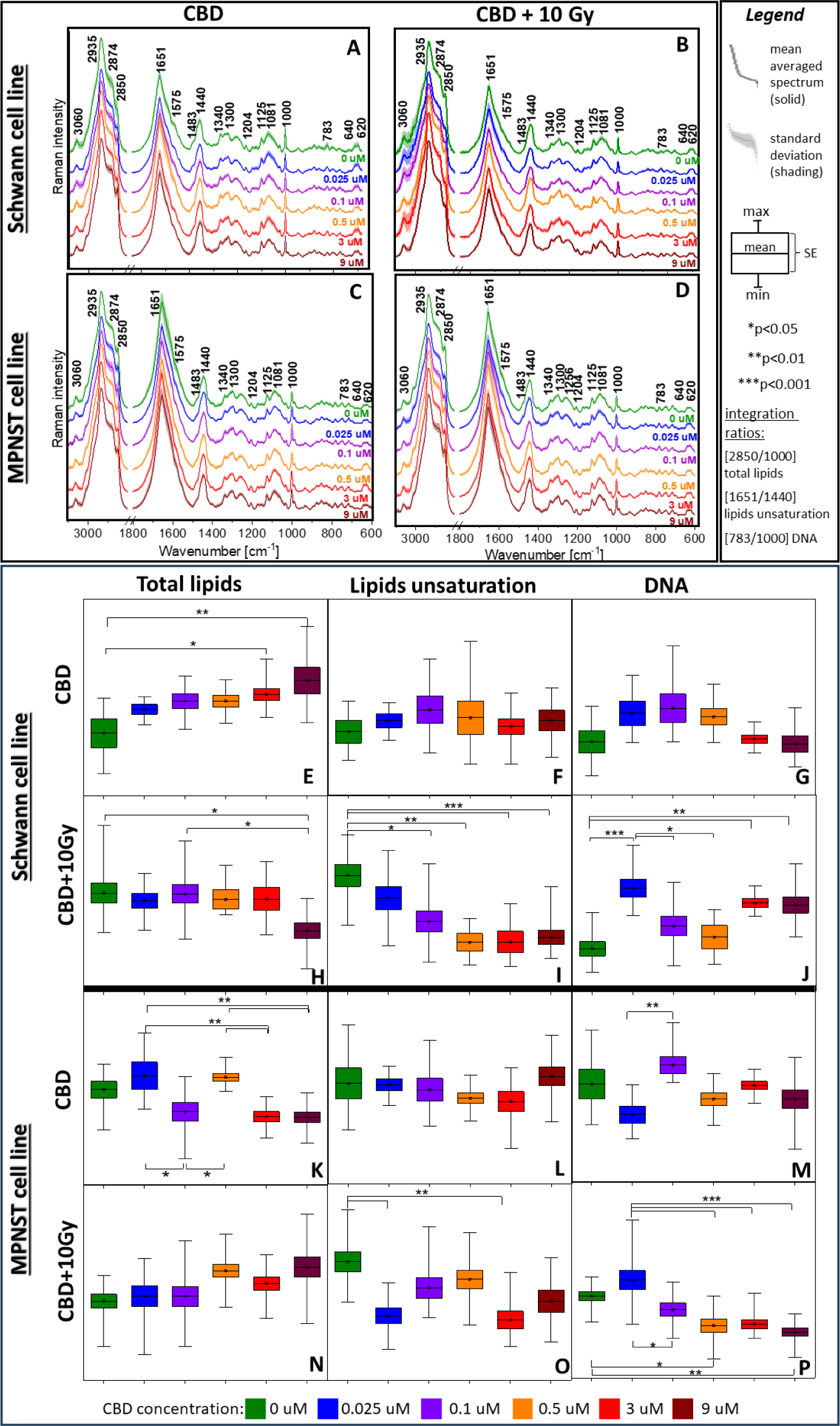
<u>Top panel:</u> Mean averaged Raman spectra from Schwann and MPNST whole cell area (A-D) incubated with CBD (A, C) and combination of CBD with 10Gy X-ray exposure dose (B,D). <u>Bottom panel:</u> Box diagrams of integral intensities for the selected Raman bands showing biochemical changes in Schwann and MPNST cells induced by CBD (E-G,K-M) and CBD with 10Gy X-ray exposure dose (H-J, N-P). The bands’ assignment is given in Table 1 (supporting information).

CBD itself induces changes in the amount of total lipids for both cell lines. An increase in total lipids concentration for Schwann cells can be related to CBD indirect enhancement of the production of long-chain polyunsaturated fatty acids such as e.g. endocannabinoids, or increased lipid storage (Fig. 4E).^50, 51^ Interestingly, the in case of cancer cells, where lipid synthesis is usually increased, CBD possibly inhibits this process (MPNST, Fig. 4K). without influencing lipid unsaturation for both cell lines (Fig. 4 F, L).^52^ Raman spectroscopy did not indicate any relevant changes in the DNA of both cell lines, influenced by CBD itself what is consistent with the comet results for t-DNA_0Gy_ (Fig. 4 G, M).

By contrast, after the application of CBD combined with radiation, for Schwann cells the total amount of lipids significantly decrease and reaches the lowest concentration for 9µM CBD, while in cancer cells the opposing trend is observed, but is not statistically relevant (Fig. 4 H, P). In case of lipids unsaturation, after radiation their amount decreases for both cell lines (Fig. 4 I, O) however for Schwann cells, this change is more gradual than for MPNST. According to the literature, the decrease in total lipids as well as their unsaturation may be related to their oxidation caused by ROS generated from ionizing radiation, which induce cell stress and lipid peroxidation.^53, 54^ For MPNST the increase in lipids relates to stress during which they release fatty acids from stored lipid droplets in order to possess energy for cell survival and repair.^55^

In the case of DNA, after irradiation, as for the total lipids, the adverse trend of changes is observed for each cell line (Fig. 4 J, R). Interestingly, Schwann cells present a significant increase in the nucleic acid concentration for the 0.025μM, 3 μM and 9 μM CBD. As CBD exhibit neuroprotective and antioxidant properties, it might decrease the concentration of free radicals and their harmful effects on DNA for normal cells.^56^ On the contrary DNA amount decreases for MPNST cells starting from 0.1 μM CBD, reaching the lowest values for the concentration of 3 and 9 μM. Therefore, with the increased CBD concentration the cancer cells become more vulnerable to radiation therapy, thus it acts as a radiosensitizer and increase the DNA damage.^57^ The observed changes for both cell lines coincide with the performed comet assay (Fig. 3 D), which determines decreased sensitivity in Schwann cells and increased sensitivity in MPNST cells to ionizing radiation upon CBD treatment in the dose dependent manner.

Raman spectroscopy was useful for the assessment of CBD and radiation influence on lipids and their unsaturation and alterations in DNA. However, to obtain profound information about protein transformation, modification in secondary protein structure as well as cholesteryl esters or molecules as phospholipids and carbohydrates, FT-IR imaging was introduced. The integral intensities calculations for mean second-derivative FTIR spectra obtained from the whole cell area allow to perform semi-quantitative analysis and provide information about the behavior of above mentioned molecules (Fig. 5).

**Figure 5.**
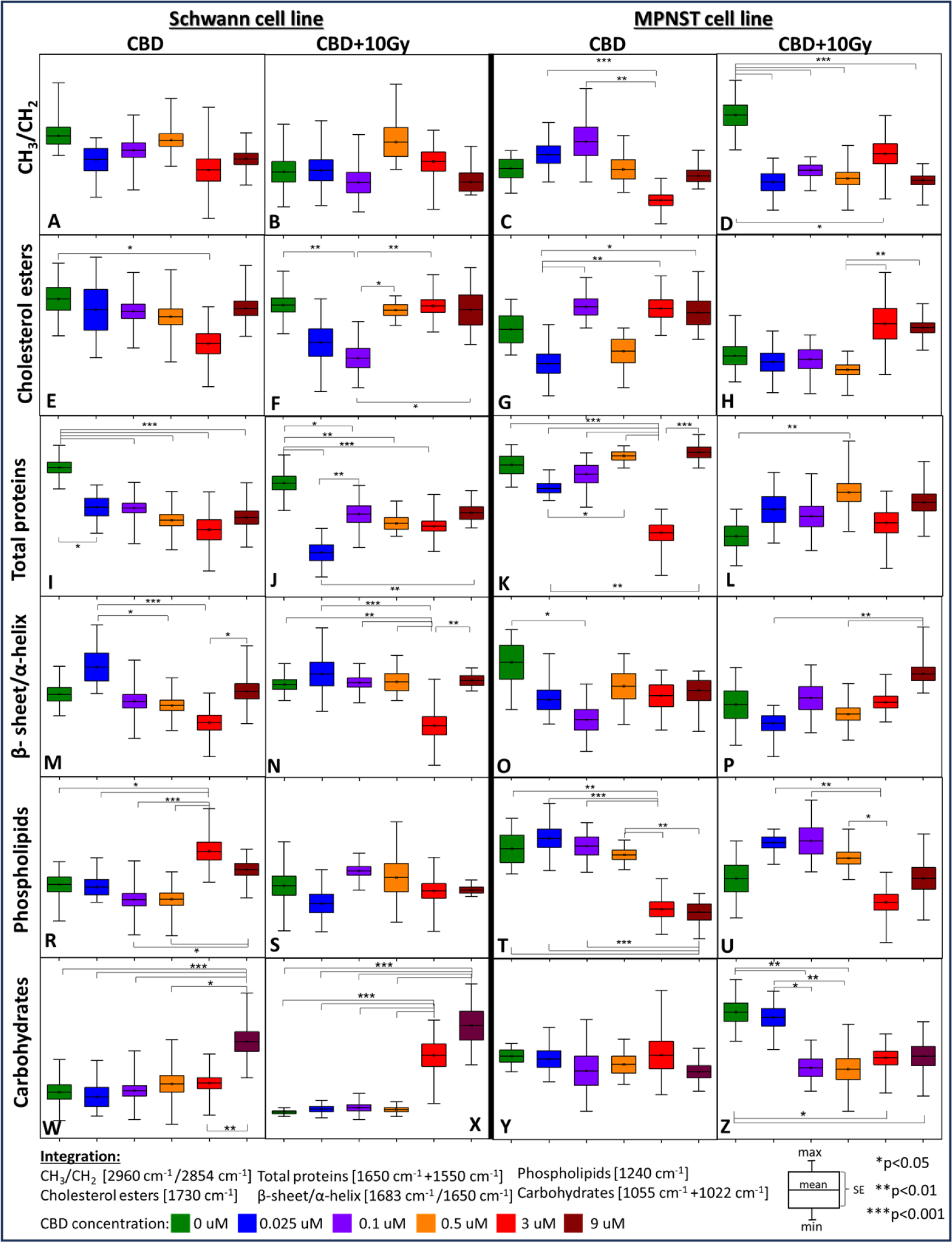
Comparison of box diagrams of integral intensities for the selected FT-IR bands showing biochemical changes in Schwann and MPNST cell lines induced by CBD and its combination with 10Gy X-ray exposure dose (A-Z). The bands’ assignment is given in Table 2 (supporting information).

The ratio of CH_3_ to CH_2_ vibrations represents the alterations in the length of fatty acyl chains, however in Schwann cells both CBD as well as its combination with X-rays did not affect the chain length (Fig. 5 A, B), where Raman indicate the decrease in lipids unsaturation after radiation (Fig. 4I) ^58^. Therefore, after radiation, with increasing CBD concentration the fatty acids chain lengths are not disturbed in Schwann cells, but become more saturated and become important source of cell energy.^59^ For MPNST treated with CBD, the observed transformations in fatty acyl chains appears for 0.1 μM, 0.25 μM, in comparison to 3 μM however are not significant in contrary to untreated cells. This also coincide with no changes in lipids unsaturation, based on Raman results (Fig.4L). Interestingly, after radiation, a strong decrease in CH_3_/CH_2_ ratio results in the predominance of -CH_2_ moieties. Such increase in -CH_2_ moieties were noticed in cancer cells after radiotherapy, and were interpreted as factor predicting apoptosis.^60^ As increase level of –CH_2_ groups correlates to apoptosis, such observation coincide with decrease DNA related to DNA damage after radiation (Fig. 4R) thereby with increase radiosusceptibility (Fig. 3D) for MPNST indicated by comet assay result.

The cholesteryl esters fraction is affected in both cell lines, interestingly for Schwann cells a significant decrease in the amount of these molecules only for 3 µM CBD concentration, in contrary to untreated cells (Fig. 5 E). It is known that CBD decrease the amount of fatty acids (FA), from which during esterification the cholesteryl esters are synthesized, therefore it is possible that for 3 µM CBD the FA are less accessible therefor lower amount of cholesteryl esters is present.^61^ Combining CBD with irradiation firstly, the decrease in cholesteryl esters for the 0.1 µM CBD dose is observed, than increase in this molecules to the level present for untreated cells from 0.5 µM till 9µM of CBD concentration. This distinct behavior of cholesteryl esters under the influence of CBD and X-rays suggests different pathways of cholesteryl esters metabolism. In case of cancer cells, CBD itself generates distinct behavior for cholesteryl esters than for Schwann cells, where only 3µM concentration significantly decrease their amount. For MPNST various fluctuation in the level of cholesteryl esters between diverse CBD concentrations is present, however not related with untreated cells (Fig. 5 G). After irradiation, the increased level of cholesteryl esters was more prominent for higher CBD concentrations (3 µM and 9µM), as displayed for Schwann cells (Fig. 5 H). These might indicates that CBD increase the cholesterol biosynthesis, as well as increase storage cell in cells for both, cholesterol and cholesteryl esters.^61^

Looking at the protein levels in Schwann cells it can be observed that CBD treatment cause total protein to decrease as CBD concentration increase. This effect is partially preserved after X-ray treatment (Fig. 5 I, J). The opposite tendency is present for cancer cells, where the protein content decreases dramatically only for the 3µM CBD concentration. Combining CBD treatment with irradiation display similar trend in the level of proteins as before irradiation, however, it is less manifested for MPNST (Fig. 5 K, L). Because CBD itself affect proteins by various pathways, mainly through protein inhibition or activation, the spectroscopic response may differ between Schwann and MPNST cell lines.^63^

Besides the determination of changes in protein amount, it was also possible to investigate the contribution of secondary protein structures. In Schwann cells, the 0.025µM CBD directs the protein structure towards β-sheet, however as the CBD concentration increases α-helical proteins begin to predominate and display the greatest contribution for 3µM CBD concentration. After irradiation, the presence of the β-sheet is no longer distinctive, but for 3µM the existence of the α-helix is still strongly manifested (Fig. 5 M, N). The appearance of β-sheet protein usually is not desirable due to its possible aggregation and transformation to amyloid present neurodegenerative diseases or in cancer.^39, 64^ Interestingly, higher CBD concentration might modify the protein structure from β to α-helix or even suppress the formation of β-amyloids.^65^ For MPNST, CBD itself generates highest amount of α-helical proteins structures, and at higher concentrations the level of β-sheet/α-helix remains at the same amount as for untreated cells. Interestingly, after irradiation, the contribution of β-sheet increases with CBD concentration and is the highest for 9µM (Fig. 5 O, P).

In the case of CBD treatment alone for Schwann cells it already causes an increase in content of the phospholipids at high concentrations (3 µM and 9µM), whereas irradiation does not change their content compared to the control level (Fig. 5 R, S). The CBD might increase the phospholipids fraction, due to direct interaction with these molecules, therefore after radiation their level was not affected.^66^ In turn, for cancer cells at the same CBD concentrations (3µM and 9µM), their content decreases significantly after treatment, opposing to Schwann cells, and this trend persists after irradiation (Fig. 5 T, U). Even though phospholipids increase in cancer cells membrane, due to disturbed metabolism, CBD reduce their amount and possibly facilitate easier cell damage with the ionizing radiation. ^67, 68^

CBD induced changes in carbohydrates level occurred to be significant for Schwann cells, where increased level of such molecules was especially observed for 9µM CBD dose compared to other concentrations and persist after X-ray irradiation additionally also for 3µM (Fig. 5 W, X). In turn, CBD itself does not cause any changes in the carbohydrate fraction of cancer cells, but in combination with radiation it significantly reduces their content even at a concentration as low as 0.1µM. Carbohydrates not only play essential role as an energetic source but also in recovery process.^69, 70^ As observed for Schwann cell line this process is more effective for higher CBD concentrations, however in the case of cancer cells it is disturbed and may contribute to increase radiosensitivity.

### 3.3 Does global FT-IR information correspond to local nanospectroscopic spectral signature?

As previously mentioned, the discussed above FTIR data were calculated from mean second derivatives spectra, obtained from whole cell area. Conventional FT-IR system imaging allows to assess rather bulk spectroscopic information due to its advantage in rapid scan collection which provides general information from a statistical number of studied samples, with the offered spatial resolution of 1.1µM. Unfortunately, such resolution is not enough for deeper insight into changes that may occur locally in cells and are related to the morphology and spatial arrangement of biomolecules. Therefore, for a better understanding of the modifications induced in cells by CBD and radiation, AFM-IR nanospectroscopy was introduced. Due to the nanometer sizes of the AFM-IR tip, it was possible to collect spectra and chemical images with the spatial resolution of 40 nm.

As can be seen in Figure 6, AFM-IR second derivatives spectra differ not only between the Schwann and MPNST cell lines but also subsequently change with increase CBD treatment. Therefore, this technique reveals more detailed biochemical information than conventional FTIR, for which the presented spectra are very similar.

**Figure 6.**
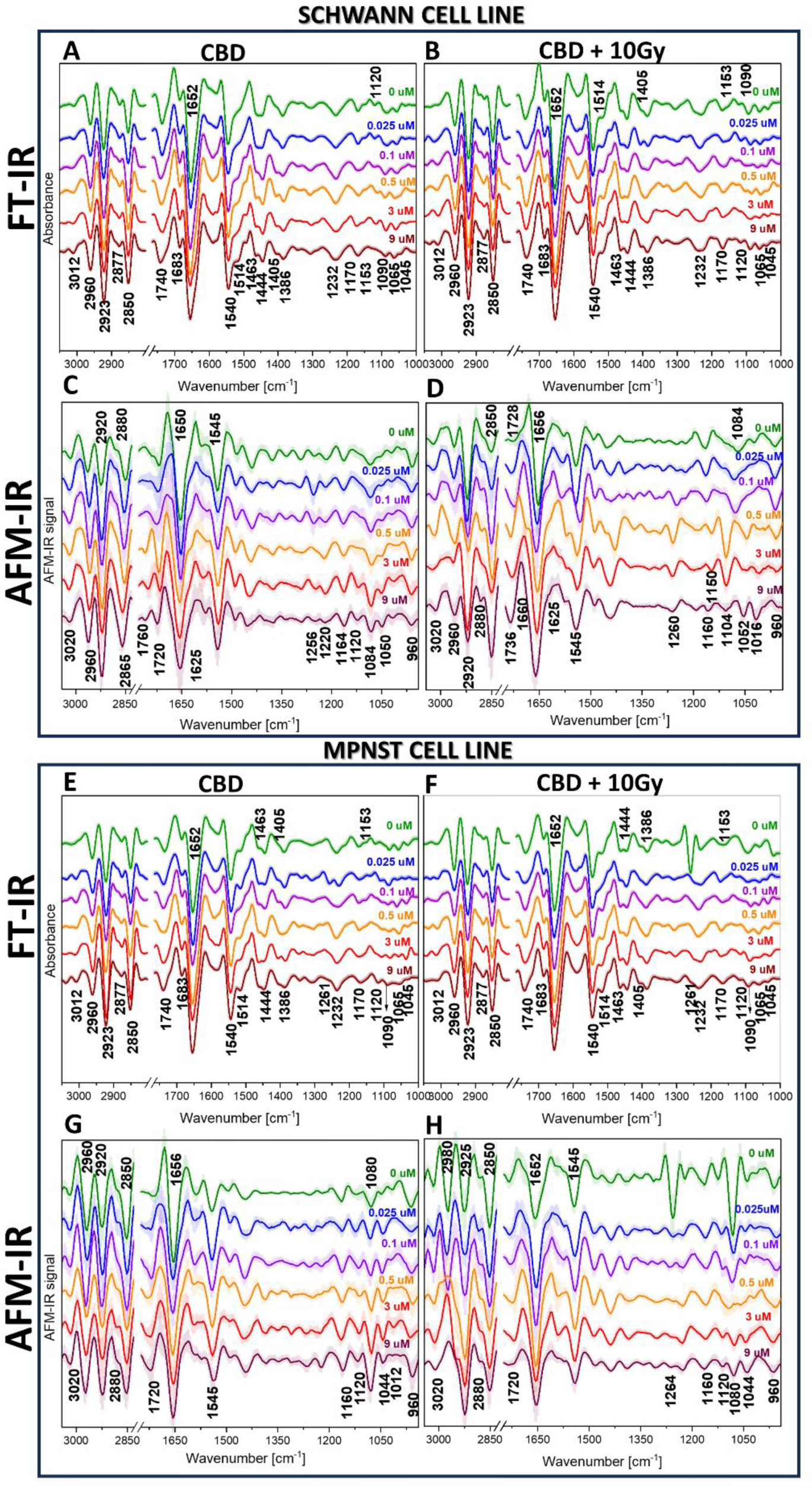
Correlation of mean second derivative spectra collected for Schwann and MPNST cell lines incubated with CBD and CBD with 10Gy X-ray exposure dose: for conventional FT-IR (A,B,E,F) and AFM-IR system (C,D,G,H). Shading denotes standard deviation. The bands’ assignment is given in Table S2 (supporting information)

For CBD-treated Schwann cells, AFM-IR revealed significant intensity changes in the bands related not only to the cholesteryl esters (1760 cm^-1^) but also fatty acids (1720 cm^-1^) indicating the alterations in this lipids fraction (Fig. 6 C). Surprisingly, band characteristic for phosphates (1232 cm^-1^) splits for two separate signals ca. 1256 cm^-1^ and 1220 cm^-1^, what might indicate the changes in the conformation of the molecule-containing phosphates e.g., in phospholipids or phosphoproteins. Also, a band related to molecules with C-O groups (1120 cm^-1^) is not present for control and 0.025µM CBD concentration however it increases gradually from 0.1µM to 9µM, therefore CBD impact the ribose (RNA) and polysaccharides in dose dependent manner. In contrary to conventional IR (Fig. 6 A), RNA-related bands are present for all CBD concentrations excluding only the highest 9µM dose. Also, fluctuation in the 1080 cm^-1^ (DNA) band intensity is more prominent for AFM-IR than conventional FTIR. In the case of irradiated Schwann cells, AFM-IR displays a significant shift from 1728 cm^-1^ (untreated cells) to 1736 cm^-1^ (9µM CBD) band (Fig. 6 D). Therefore, CBD not only leads to the alterations in cholesteryl esters amount according to previously showed box plots (Fig. 5 E-H) but also their structure. Interestingly, in contrast to non-radiated Schwann cells, for 0.5µM and 3µM concentrations strong band at 1260 cm^-1^, 1150 cm^-1^ and a significant shift from 1084 cm^-1^ related to phosphates in DNA and phospholipids to 1104 cm^-1^ is presented, but this trend disappears for 9µM CBD dose. Described changes in DNA conformation which might result in radioresistence mechanism.

For cancer cells incubated with CBD, FT-IR show similar lipid profiles between the conditions (Fig. 6E), while AFM-IR discriminates subsequent alterations in the 3020 cm^-1^ -2850 cm^-1^ spectral range (lipids) mainly changes in 2960 cm^-1^ to 2920 cm^-1^ bands ratio.

Interestingly, in contrary to FTIR (Fig. 6 E) AFM-IR did not reveal the presence of 1683 cm^-1^band, related to β-sheet protein structure, therefore this protein conformation aggregates might be deposited locally within investigated cells, what was visible for overall cells screening by FTIR. More prominent changes occur in 1160 cm^-1^ -1012 cm^-1^ spectral region, where the band 1080 cm^-1^ intensity increase gradually with CBD concentration (Fig. 6G). This might relate to increase DNA activity. Since increased production of DNA is not possible, we hypothesize that DNA can accumulate locally, and this can be detected by AFM-IR.

After radiation the most significant changes are visible for MPNST cells without CBD treatment, manifested by intense band at 1261 cm^-1^ what is observed for both FTIR and AFM-IR spectral profile (Fig. 6 F, H). For AFM-IR spectra this band is more prominent and additionally accompanied by sharp 1080 cm^-1^ band, which decrease significantly with the increase CBD concentration. This is possibly the result of DNA damage and decay of phospholipids resulting from ionizing-radiation, however the presence of CBD weakens X-rays toxicity, therefore this bands were not observed for higher concentrations.^71, 72^ Also the lipid profile of the MPNST cells changes dramatically, however only for irradiated cells treated with CBD. Such alterations can be seen only by AFM-IR technique as the 2980 cm^-1^ band shifts towards 2925 cm^-1^ and appears as a shoulder for 3μM CBD dose (Fig. 6 H). In contrary, the profile of 2980-2850 cm^-1^ range for the irradiated but not treated with CBD MPNST cells display the same pattern as for untreated MPNST cells before irradiation (Fig. 6G). Based on these observations, the alterations in lipids present for the irradiated cancer cells are the result of the CBD treatment combination with radiotherapy, hence not implicate from irradiation itself. Consequently, this alterations in lipids may play key role in the increase radiosuscebility for MPNST cells.

Finally, comparing the spectral information between two infrared spectroscopic techniques, for Schwann and MPNST cells, at the first glance the FTIR spectra are similar for both treated with CBD as well as for CBD combined with irradiation (Fig. 6 A, B, E, F). Only noticeable changes are related to cancer cells with 0.025µM CBD dose (Fig. 6E) where the 1261 cm^-1^ and 1232 cm^-1^ bands occurred, in contrary to single band at 1232 cm^-1^ for Schwann cells treated the same dose of CBD. AFM-IR technique reviled visible changes in the 2980-2850 cm^-1^, where the changes in 2960 cm^-1^ to 2920 cm^-1^ ratio is characteristic for each cell line and spectral pattern is enhanced gradually with increase CBD concentration (Fig. 6C, G). After X-ray exposure changes between normal and cancer cells indicated by FT-IR are observed mainly for untreated cells, where after irradiation MPNST display strong band at 1261 cm^-1^, absent for untreated and irradiated Schwann cells. This band is also distinguishable with AFM-IR technique, however is more prominent and for cancer cells beside 1264 cm^-1^ band additional 1080 cm^-1^ band is observed (Fig.6 H). Such spectral pattern is not observed for untreated, irradiated Schwann cells (Fig. 6D). Other significant alterations revealed by AFM-IR for irradiated cells are related to lipids for both cell lines, especially in the range of 0.5µM to 9µM CBD dose. (Fig. 6 D, H) This shows various modification in lipid metabolism induced by radiation. Moreover, also for these concentrations the different phosphate response is present e.g., for Schwann intense 1260 cm^-1^ band is present and significant band shift (1084 cm^-1^ -1104 cm^-1^) is observed what is not visible for cancer cells. Interestingly, AFM-IR indicates the presence of β-sheet proteins only for Schwann cells (1625 cm^-1^) before and after irradiation.

### 3.4 AFM-IR chemical maps reveal changes in the distribution of selected molecules

So far, the discussed results based on Raman and FT-IR spectral profiles allowed for the determination of molecules and semi-quantitative analysis of the level of alterations resulting from the CBD and irradiation effect for Schwann and MPNST cell lines. The AFM-IR technique provide information with the possible changes in the orientation or structure of investigated molecules. To fulfill this analysis, also the AFM-IR maps were collected for topography and chemical images for selected molecules.

Because biological tests and spectroscopy studies reveal that from 3µM concentration most significant changes related to CBD treatment occurred, exemplary nanoscopic infrared maps with the 40nm spatial resolution were collected for the amide I and cholesteryl esters band to evaluate the local changes for these molecules for Schwann cells (Fig. 7) and for MPNST cells (Fig. 8).

**Figure 7.**
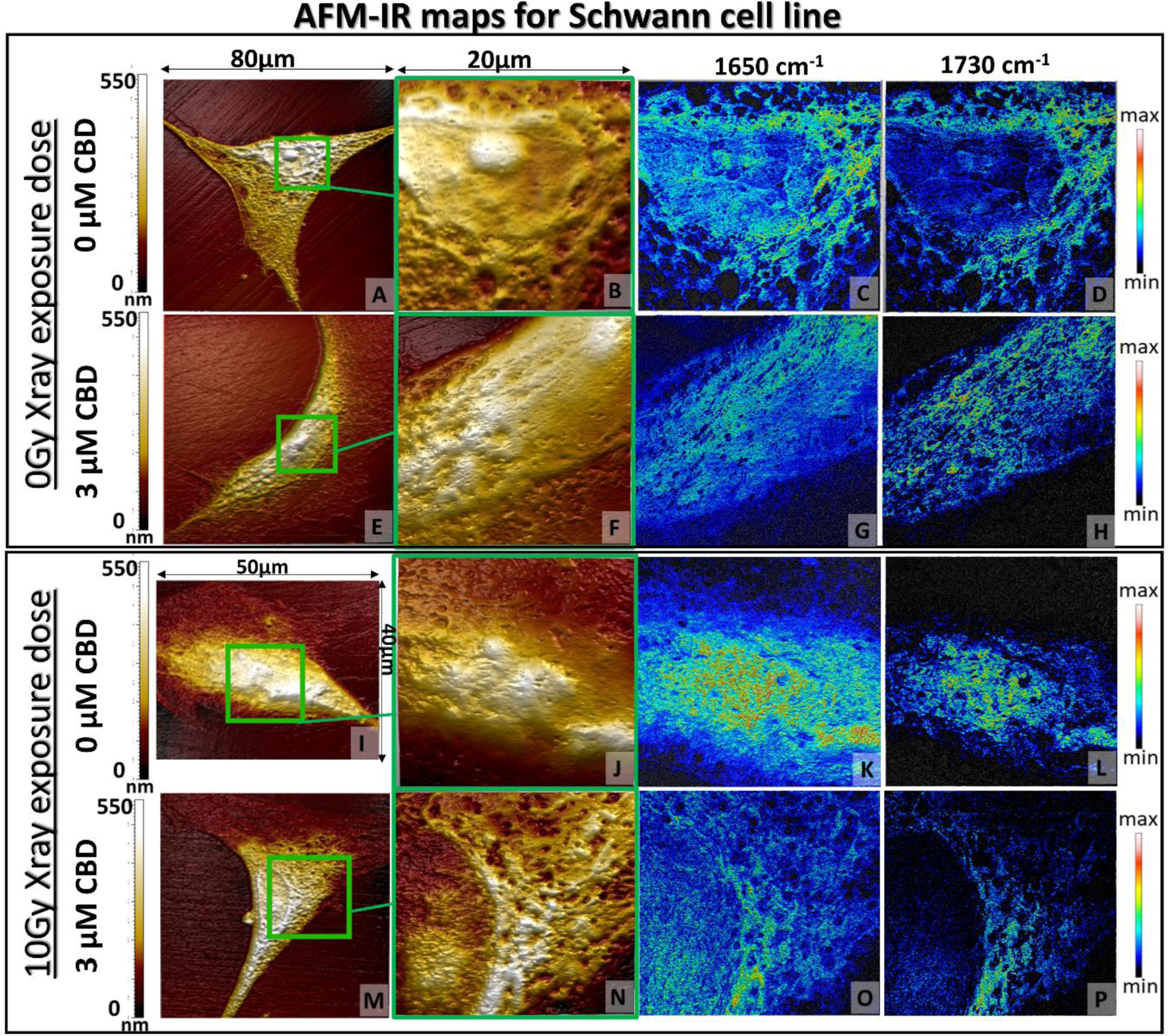
AFM topography of the whole Schwann cell (A,E,I,M) and selected area (green box) of 20×20µM (B,F,J,N) for nanoscale chemical mapping at 1650 cm^-1^ and 1730 cm^-1^ bands (C,D,G,H,K,L,O,P). Collected images were selected for 3µM CBD concentration in reference to the CBD untreated cells before (top panel) and after 10Gy X-ray dose application (bottom panel).

**Figure 8.**
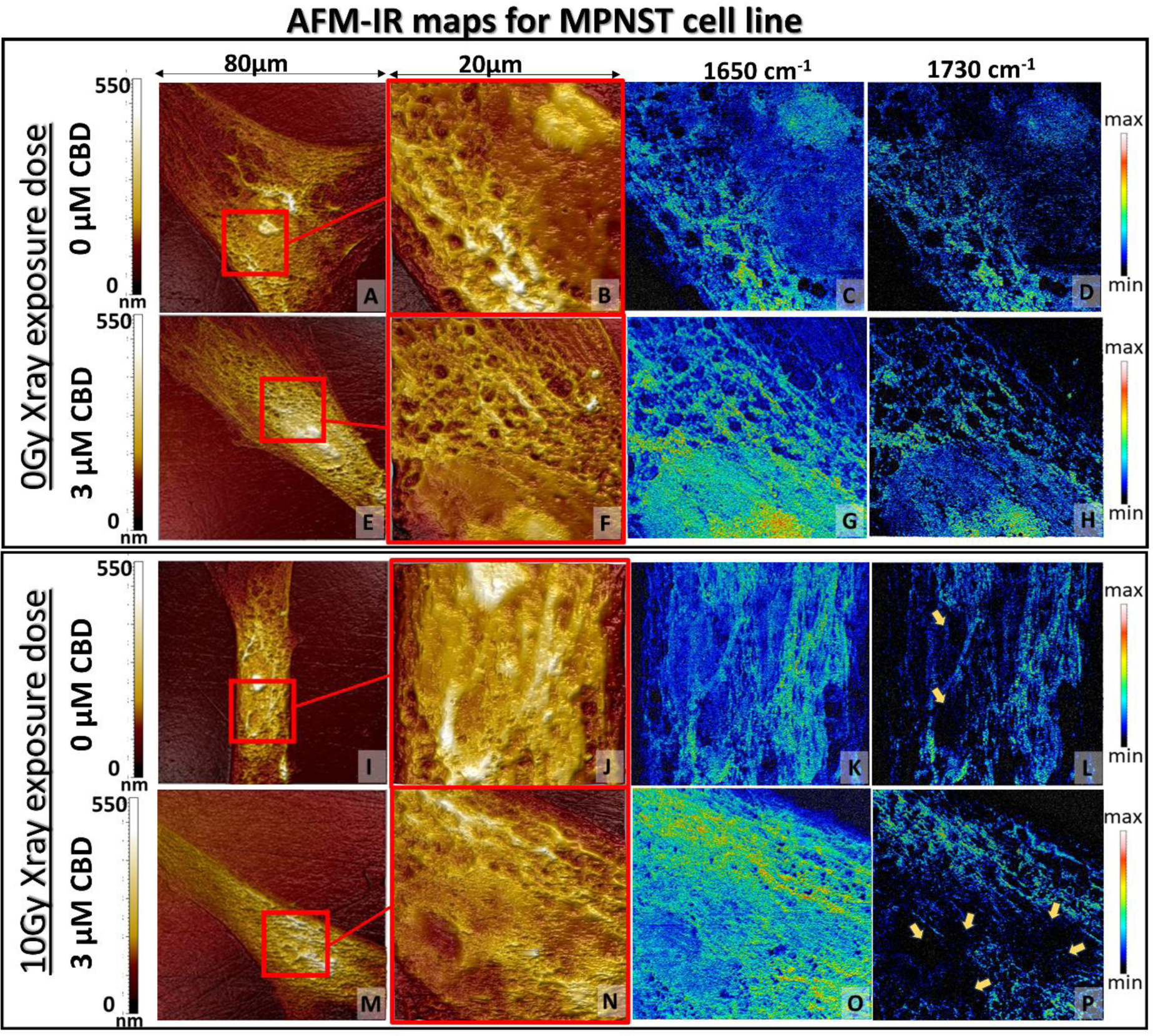
AFM topography of the whole MPNST cell (A,E,I,M) and selected area (red box) of 20×20µM (B,F,J,N) for nanoscale chemical mapping at 1650 cm^-1^ and 1730 cm^-1^ bands (C,D,G,H,K,L,O,P). Collected images were selected for 3µM CBD concentration in reference to the untreated cells without (top panel) or with 10Gy X-ray dose application (bottom panel).

Non-treated and non-radiated Schwann cell lines morphology present cell morphology, with a with a well-defined nucleus and nuclei, surrounded by cytoplasm containing pores can be seen at Fig. 7 A and B. An AFM-IR chemical map of the Schwann cell imaged at 1650 cm^-1^ (Fig. 7 C) relates to proteins, which are distributed evenly within whole cell area and correlate with the cell morphology. On the contrary, the 1730 cm^-1^ spectral signal corresponding to the cholesteryl ester is localized mainly around the nucleus, probably in its endoplasmic reticulum. ^73^ After the incubation with 3µM concentration of CBD, the height image of the cell area present also clear distribution of the cell component such as nuclei sub-nuclear area, however the margins between each subcellular elements are less visible (Fig. 7 E, F). The protein distribution overlays with the cholesteryl esters distribution and do not localize in particular cell area such as it was observed for untreated cell (Fig. 7 G, H). The use of 10Gy X-ray exposure dose on Schwann cells without CBD cause alterations in the cell morphology (Fig. 7 I, J). The nuclei and cytoplasm have no clear margins therefore it is difficult to determine their localization. Also, the protein distribution is concentrically accumulated and as previously described corresponds to the cholesteryl esters distribution (Fig. 7 K, L). Surprisingly, the cells incubated with 3µM CBD concentration without radiation allowed to preserve the normal cell morphology, with clearly differentiative nucleolus, nuclei, and its membrane as well as cytoplasm as for control (non-treated and not irradiated) cells (Fig. 7M, N). Also, the chemical maps shows a more localized distribution, cholesteryl esters are gathered around the nuclei and do not overlay with the protein signal (Fig. 7O, P).

AFM topography images of MPNST cells not exposed to X-ray radiation are similar, and the 3µM CBD concentration did not change their morphology (Fig. 8 A,B,E,F). They both exhibit a clearly outlined nucleolus within the nucleus separated by a membrane from the cytoplasm. In the case of protein and cholesteryl esters distribution, the more intense signal from these molecules is observed for cells treated with 3µM CBD concentration, interestingly also in the nucleolus and nucleus area (Fig. 8 C, D, G, H). When cancer cells were exposed to 10Gy X-ray dose, significant changes in the cell morphology appeared. That manifested mainly in the nuclei deformation for both 0µM and 3µM of CBD concentrations however, the nucleolus was still differentiable for 0µM CBD concentration (Fig. 8 I, J, M, N). The combination of 3µM CBD with X-rays induced the disappearance of the nuclei membrane and numerous hollows in the nuclei area. For both, irradiated and non-irradiated cells the protein signal is distributed within the whole cell area however is more prominent with 3µM CBD treatment. The localization of cholesteryl esters is disturbed for both irradiated cells without and with CBD treatment. The 1730 cm^-1^ signal is scattered within the whole cell area and is absent in areas corresponding to hollows presented by AFM topography (Fig. 8 L, P yellow arrows) which corresponds to the nuclei deformation.

The most significant changes in the morphology and IR chemical maps are observed for Schwann and MPNST cells after radiation (Fig. 7 and 8 I-P). They prove that CBD treatment play a key role in the cell morphology modification in both cell lines, however in the different manner. For Schwann cells, despite the ionizing radiation, CBD treatment preserves normal morphology and protein and cholesteryl esters distribution, whereas for MPNST CBD strengthens the ionizing radiation harmful effect which manifests in alterations in both morphology and biomolecules distribution on chemoselective maps.

## 4. Conclusions

Performed studies indicate that the cells radiosensitivity can be modified under the influence of CBD, which increases the toxicity of ionizing radiation in MPNST cells and simultaneously decrease it for Schwann cells. The use of hyperspectral imaging and AFM-IR allowed to broaden the knowledge in the field of biochemical modification introduced by CBD in the peripheral nervous system *in vitro* model. The CBD targets different molecules and exhibits different mechanism of action when combined with X-ray radiation for normal and cancerous cell line.

The most significant alterations indicated with Raman spectroscopy were in the total amount of lipids and DNA, accompanied with changes reviled by AFM-IR in lipid bands ratio and shifts for DNA band, which may contribute to DNA conformation changes after irradiation of Schwann cells treated with CBD. Other molecules, which were modulated under the influence of CBD and ionizing radiation, suggested by FT-IR and AFM-IR, were cholesteryl esters and phospholipids. FT-IR display the significant changes in their concentration depending on applied treatment and cell line. Nanospectroscopy provides information about the possible modification in the conformation of molecule-containing phosphates and cholesteryl esters indicated by strong band shifts, which may play a key role in the decreased radiosensitivity for Schwann cells. Besides, AFM-IR chemical maps display the most prominent alterations in the protein and cholesteryl esters distribution for both cell lines, after the CBD incubation and X-ray radiation. Schwann cells preserve similar distribution of these molecules after tretment, however in MPNST cells CBD triggers the alterations in their localization as well as in the cell morphology.

## 5. Acknowledgments

K. Chrabaszcz thank the National Science Centre, Poland (MINIATURA, 2022/06/X/ST4/00414) for financial support of the research. The study was performed using equipment purchased in the frame of the project co-funded by the Malopolska Regional Operational Program Measure 5.1 Krakow Metropolitan Area as an important hub of the European Research Area for 2007– 2013, Project No. MRPO.05.01.00-12-013/15.

