## Supporting information for "High resolution optical spectroscopy for the evaluation of cannabidiol efficiency as a radiation therapy support of peripheral nervous system tumors"

**Table S1.** Raman band positions observed in Raman spectra of the Schwann and MPNST cell lines with their assignment to biomolecules.[1-5]

| POSITION [$CM^{-1}$] | ASSIGNMENT TO BIOMOLECULES AND VIBRATIONAL MODES |
|---|---|
| **3060** | Unsaturated fatty acids; ν(=C-H) |
| **2936** | Lipids, proteins; ν(C-H) |
| **2874** | Lipids, proteins; ν(C-H)-$CH_2$ |
| **2850** | Long chain fatty acids; $\nu_s(CH_2)$ |
| **1651** | Proteins (amide I); ν(C=O) and δ(N-H) |
| **1575** | Tyrosine |
| **1440** | Proteins, lipids; δ(CH2), δ($CH_3$) |
| **1340** | A (nucleic acids); δ(CH)<br>Proteins; δ(CH)<br>Carbohydrates; δ(CH) |
| **1300** | Lipids; τ$CH_2$-$CH_3$ |
| **1204** | Tyrosine, Phenyloalanine, Tryptophan, Hydroxyproline (proteins); τ($CH_2$) |
| **1125** | Lipids (trans in acyl backbone); ν(C-C)<br>Cyt; ν(C-N)<br>Proteins; ν(C-O)<br>Carbohydrates; ν(C-O) |
| **1081** | Phospholipids; ν($PO_2^-$)<br>Lipids (trans in acyl backbone); ν(C-C) |
| **1000** | Phenylalanine (symmetric ring breathing) |
| **783** | DNA; $\nu_{as}$(O-P-O)<br>Uracyl, Thymine, Cytosine (nucleic acids) |
| **640** | Tyrosine |
| **620** | Phenylalanine |

ν – stretching mode, as – asymmetric, s – symmetric; δ – in-plane deformations; τ – twisting;

**Table S2.** IR band positions observed in second derivatives infrared spectra of the Schwann and MPNST cell lines with their assignment to biomolecules. [6-11]

| POSITION [$CM^{-1}$] | ASSIGNMENT TO BIOMOLECULES AND VIBRATIONAL MODES |
|---|---|
| **3012,3020** | Unsaturated fatty acids; ν(=C-H) |
| **2960** | Proteins, lipids; $\nu_{as}(CH_3)$ |
| **2923,2920** | Lipids and proteins; $\nu_{as}(CH_2)$ |
| **2877** | Proteins, lipids, nucleic acids; $\nu_s(CH_3)$ |
| **2880** | Terminal CH3 group in acyl chains (lipids); ν(CH) |
| **2850/2866** | Long chain fatty acids; $\nu_s(CH_2)$ |
| **1740** | Triacylglycerols; $\nu_{ester}$(C=O) |
| **1728** | Cholesterol esters; $\nu_{ester}$(C=O) |
| **1720** | Fatty acids; ν(C=O)<br>Base pair (B-DNA); ν(C=O) |
| **1683** | β-turns in proteins (amide I); ν(C=O) and δ(N-H)<br>Guanine (DNA); ν(C=O) and ν(C=C) |
| **1660-1650** | α-Helices in proteins (amide I); ν(C=O) and δ(N-H) |
| **1625** | β-sheet in proteins (amide I); ν(C=O) and δ(N-H) |
| **1540-1545** | Proteins (amide II); δ(N-H) and ν(C-N) |
| **1514** | Tyrosine (proteins); ν(CC) of the Tyrosine ring<br>Cytosine (methylated DNA); in-plane vibrations of the ring |
| **1463** | Proteins; δ($CH_2$, $CH_3$)<br>Cytosine (DNA); δ(NH), ν(CC) |
| **1444** | Lipids; δ($CH_2$, $CH_3$) |
| **1386** | Free fatty acids; $\nu_s(COO^-)$<br>Free amino acids; $\nu_s(COO^-)$ |
| **1260-1220** | Nucleic acids, phospholipids, phosphoproteins; $\nu_{as}(PO_2^-)$ |
| **1170-1164** | Fatty acids and cholesterol esters; ν(C-O) |
| **1150-1153** | Glycogen; $\nu_{as}$(CO-O-C)<br>Polysaccharides; $\nu_{as}$(CO-O-C) |
| **1120-1104** | Ribose (RNA); ν(C-O)<br>Polysaccharides; ν(CC-OC) |
| **1090-1084** | Nucleic acids; $\nu_s(PO_2^-)$<br>Phospholipids; $\nu_s(PO_2^-)$<br>Glycogen; ν(C-C) |
| **1065-1044** | Carbohydrates, glycogen; ν(C–O) |
| **960** | DNA; ν(C-C) |

ν – stretching mode, as – asymmetric, s – symmetric; δ – in-plane deformations;
